# Genome-wide mapping of sex-associated autosomal DNA methylation in major blood cell types

**DOI:** 10.1101/2025.10.14.682299

**Authors:** Yutaro Yanagida, Yutaka Nakachi, Miki Bundo, Syohei Komaki, Atsushi Shimizu, Kazuya Iwamoto

## Abstract

Sex differences shape human physiology and disease risk, yet their autosomal epigenetic basis remains incompletely understood. Using whole genome bisulfite sequencing data from iMETHYL database (∼100 adults) across three purified blood cell types (CD4⁺ T cells, monocytes, neutrophils), we performed comprehensive mapping of cell-type specific sex-associated DNA methylation. Using genome-wide *Z*-test and confidence-interval method, we identified thousands of autosomal differentially methylated sites (DMSs), the majority of which were cell type-specific. We found that neutrophils exhibited pronounced female-biased hypermethylation, whereas CD4⁺ T cells and monocytes showed more balanced patterns. DMSs were enriched within gene bodies and annotated to neuronal and adhesion functions, with T-cell activation uniquely associated in CD4⁺ T cells. Transcription factor binding site enrichment indicated hematopoietic regulators, suggesting that sex-associated methylation is tightly linked to early developmental processes within blood lineages. Differentially methylated regions overlapped genome-wide association study loci for lifetime smoking, multiple sclerosis, and psoriasis, indicating convergence between sex-associated epigenetic states and genetic susceptibility. Evolutionary analysis revealed limited conservation but identified a conserved intronic CpG within *FIGN*. This study provides the first genome-wide, cell type-resolved map of autosomal sex-associated DNA methylation in human blood and establishes a foundation for mechanistic and translational studies in sex-informed biology and medicine.

**GRAPHICAL ABSTRACT:** 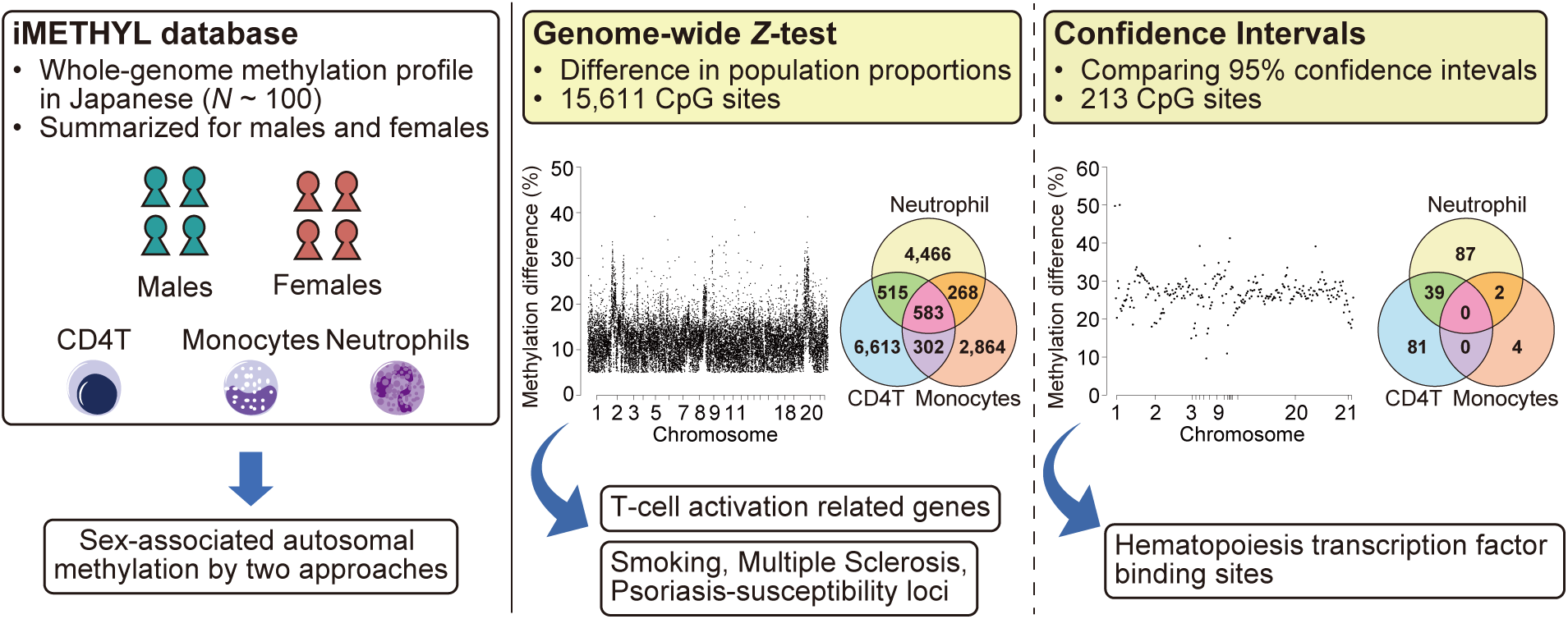

## INTRODUCTION

Biological sex profoundly influences human physiology and disease susceptibility, shaping diverse traits ranging from growth and metabolism to immune responses and neuropsychiatric outcomes (1–6). Understanding the molecular basis of sex differences is therefore essential for both basic biology and sex-informed medicine. While sex chromosomes have been regarded as the primary determinants of sex differences, accumulating evidence highlights substantial contributions from autosomes through genetic variants and expression differences (7–12).

Epigenetic modifications provide a key layer of expression regulation that may determine how autosomes drive male- or female-specific traits (13,14). Among these, DNA methylation, defined as the addition of a methyl group to cytosine in CpG dinucleotides, is a major regulatory mechanism of gene expression. DNA methylation in the promoter region typically suppresses nearby gene activity, although the effects of methylation in the gene body have been controversial (15–17). Regions enriched with CpG sites, known as CpG islands, are frequently located in promoter regions, where their methylation status is critical for tissue-specific gene expression (17,18).

Several studies have explored sex-associated DNA methylation on autosomes (19–28). A meta-analysis of 76 studies identified 184 autosomal CpGs with sex-associated methylation, 178 of which were hypermethylated in females, with a mean difference of 3.7% (19). Another study analyzing whole blood datasets reported 179 sex-associated methylated regions, with 152 hypermethylated in females. These differences were conserved across different ages and linked to sex hormone levels (20). Additionally, a study analyzing postmortem brain tissues identified 15,417 sex-differentially methylated CpGs, with significant enrichment in genes linked to psychiatric disorders (21). Another study on whole blood reported aging-related sex differences in methylation, identifying 52 autosomal CpGs including CpGs associated with traits like prostate cancer and male baldness (22). Lastly, a study identified 396 autosomal CpGs with sex-associated methylation, 293 hypermethylated in females, with functional links to genes involved in male fertility and sex development (23).

Previous studies have mainly used whole blood with commercially available microarray platforms, raising two potential limitations. Firstly, given that different cell types within blood showed different profiles, the results obtained from whole blood cells were affected by the cellular compositions (29,30), leading to false negatives of DNA methylation differences occurred in the specific cell types. Secondly, commonly used commercial microarrays can detect up to ∼850 thousand CpG sites. Given that the human genome contains approximately 30 million CpG sites (31), the vast majority (over 95%) remain unanalyzed in detail.

The iMETHYL database (30,32) consists of multi-omics dataset of peripheral blood collected from a Japanese population. It includes whole genome bisulfite sequencing (WGBS) data of distinct blood cell types (CD4+ T cells (CD4T), monocytes, and neutrophils) from approximately 100 healthy subjects, consisting of roughly equal number of males and females. We utilized the mean and standard deviation (SD) of DNA methylation levels across all autosomal CpG sites, and developed two analytical frameworks to detect differentially methylated sites (DMSs): a *P*-value approach based on the *Z* test (*P*-value-based approach) and a confidence interval (CI)-based approach with adjustable stringency. The *P*-value-based approach identified a total of 8,013, 4,017, and 5,832 autosomal DMSs for CD4T, monocytes, neutrophils, respectively. The number of unique DMSs was 15,611, the majority of which were cell type-specific. In addition, we identified 213 high-confidence DMSs by the CI-based method. These DMSs showed larger mean methylation differences, with 26.7% by the CI-based method compared to 12.1% by the *P*-value-based, and were predominantly located within gene bodies. Gene Ontology (GO) analysis revealed enrichment of neuron, synapse, and cell adhesion-related genes across all cell types, and T cell activation specifically in CD4T. Comparison with previous array-based studies showed limited overlap, likely due to differences in thresholds and genomic coverage.

Differentially methylated regions (DMRs) significantly overlapped with genome-wide association study (GWAS) loci associated with lifetime smoking status, multiple sclerosis, and psoriasis. Enrichment analysis of transcription factor (TF) binding sites identified TFs involved in hematopoiesis, suggesting that the DMSs are closely linked to blood cell differentiation. Additionally, several DMSs were found to be evolutionarily conserved across species. Our findings provide novel insights into the epigenetic landscape of sex differences in human blood cells and demonstrate the power of cell type-specific, genome-wide methylation analysis in understanding sex-biased traits and diseases.

## MATERIAL AND METHODS

### WGBS data

The iMETHYL database provides summarized WGBS data for three cell types: CD4T, monocytes, and neutrophils, each of which was isolated from whole blood samples from approximately 100 Japanese participants. Detailed methods for sample collection, WGBS, and data processing are described elsewhere (30,32). In brief, CD3+/CD4+ T cells and CD14++/CD16-monocytes were isolated from peripheral blood mononuclear cells (PBMCs) (30), while CD16+ neutrophils were isolated from polymorphonuclear cells (32) using a cell sorter. WGBS was conducted on an Illumina HiSeq 2500 or HiSeq X platform, with a read length of 125 bp or 150 bp (paired-end). Sequence reads were mapped to the GRCh37d5 reference genome using NovoAlign (ver. 3.02.08). DNA methylation levels were represented by the ratio of methylated read counts to the total read counts mapped to each CpG site. The total subject numbers for CD4T, monocytes, and neutrophils were 102, 102, and 94, respectively, with male ratio of 48.0, 47.1, and 51.1%, and mean ages of 62.0, 62.5, and 58.0 years. The sequencing depths for these three cell types were 31.0, 31.1, and 54.7, and the detected CpGs, defined as those retained in over 50% of subjects for each cell type, were 24,037,518, 23,941,821, and 25,483,031, respectively.

### Quality control (QC)

Due to the protection of privacy of participants, we used the mean and SD of DNA methylation levels at each CpG site in males and females. These data were available from the web site (http://imethyl.iwate-megabank.org) (30,32). After excluding 24 duplications of an identical CpG site in downloaded data, we excluded CpGs overlapped with the common SNPs in the Japanese population with BEDTools (33). We used SNP information in the Japanese population based on 3.5KJPNv2 reports of jMorp (34–36) for this purpose. Then, we excluded CpGs with a sample size of less than ten in males or females to maintain the quality. Among the remaining CpGs, we utilized the CpGs common to both males and females within each cell type.

### Identification of sex-associated DMS

Given the mean and SD of DNA methylation level, and sample size in males and females at every CpG site across the human genome, we constructed two different ways, *P*-value-based or CI-based approach to identify sex-associated DMS. The *P*- value-based approach consists of two-sided *Z*-test for the difference in population proportions and threshold of the actual difference of methylation level. We conducted *Z*-test with Bonferroni correction for multiple testing based on the number of all CpGs in three cell types (*N* = 69,040,758; significance level, *P*-value < 7.24 × 10^-10^). Then we retained CpGs with methylation difference 5% or more. The CI-based approach consists of comparison of CIs for methylation level in males and females and threshold of the actual difference. We firstly defined sex-associated methylation by the Wald method with the modifications by introducing two parameters, *t* and *d* as follows:

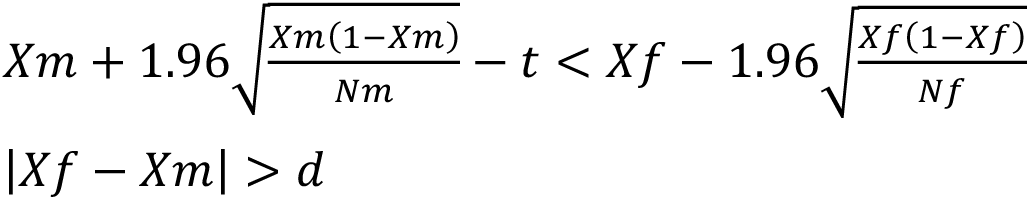

In this formula, the mean value of methylation level in males (*Xm*) or females (*Xf*), the number of males (*Nm*) or females (*Nf*) were used to identify DMS between males and females. This formula represents the case for identifying female-hypermethylated sites, and similarly applies to male-hypermethylated sites. The parameter of *t* (0 ≤ *t* ≤ 1) means the tolerance for the overlap between 95% CIs of males and females, and simultaneously indicates the difference between the upper limit and the lower limit of each 95% CI. The higher the value of *t* increases, the looser the criteria become. In addition, the parameter of *d* (0 ≤ *d* ≤ 1), which represents the threshold for differences in methylation level, was used to prevent an inflation of detected CpGs. We defined *d* as 5% a priori. We defined flanking DMSs within 1 kb as DMRs using an R package of igraph (37).

### Datasets used for comparison and interpretation of DMS

We used eight previously published datasets of sex-associated methylation using whole blood, including those from Price et al. (24), Shah et al. (25), Singmann et al. (26), Zaghlool et al. (27), Suderman et al. (28), McCarthy et al. (22), Gatev et al. (20), and Grant et al. (23). To consider the effect of smoking, we used a dataset of smoking-associated CpGs, including 972 CpGs reported by Sonja et al (38). All datasets were generated using Illumina methylation platforms. We examined the reported CpG sites and included only those that were assigned specific genomic locations in hg19 coordinates.

### Annotation and GO analysis

We conducted annotation of DMSs using an R package of annotatr (39). The genic annotation was performed regarding the regions of promoter, exon, intron, 1 to 5 kb from transcription start site, 3’UTR, 5’UTR, and intergenic regions. The CpG island annotation was performed regarding the regions of CpG islands, CpG shelves, CpG shores, and CpG inter. We conducted GO analysis using the annotated genes by ToppGene (40–42). We further performed GO analysis correcting for gene length bias by an R package of goseq (43).

### Enrichment analysis with the GWAS regions

We conducted an enrichment analysis between sex-associated DMRs and SNPs reported in previous GWASs. GWASs were selected from the GWAS Catalog database (44) (accessed on April 20, 2025) based on the following exclusion criteria: (1) unclear sample size or trait description; (2) initial sample size of fewer than 10,000; (3) studies focused on metabolite levels; (4) fewer than 1,000 initial case subjects; (5) traits described using four or more words; (6) fewer than 50 reported SNPs; and (7) studies investigating same trait but with smaller sample sizes. We used the resultant 337 GWASs. Enrichment was evaluated using odds ratios and Fisher’s exact test, based on the number of GWAS-regions overlapping or not overlapping DMRs, as well as flanking regions within 1 kb of all CpG sites in iMETHYL for each cell type. The analysis was performed using R packages of biomaRt (45,46), rtracklayer (47), GenomicRanges (48), LOLA (49), and gwasrapidd (50).

### Datasets used for gene expression analysis

We used public RNA-seq data of whole blood in GTEx Analysis V8 (51). Expression levels were compared between males and females using Mann-Whitney *U* test. The data of gene expression normalized by transcripts per million (TPM) (gene_tpm_2017-06-05_v8_whole_blood.gct.gz) used for the analyses described in this manuscript were obtained from the GTEx Portal on 06/02/2023.

### TF-binding site enrichment analysis

We evaluated the enrichment of TF-binding sites using the regions within 5 base pairs flanking the DMSs using ChIP-Atlas 3.0 (https://chip-atlas.org) (52–54), using datasets for TFs and others in blood, the threshold peak-call significance *Q*-value < 10^-5^. For each cell type and sex, background datasets were constructed by randomly selecting CpGs at ten times the number of corresponding DMSs, excluding the DMSs and their flanking regions. We also conducted a protein-protein interaction (PPI) analysis based on the ChIP-Atlas results after Bonferroni correction for multiple testing (7,397 entries × 6 tests) using stringApp (55), and visualized the network relationships with Cytoscape (56).

### Analysis of evolutionary conservation

We examined the conservation of DMSs across species using the Comparative Genomics track of the UCSC Table Browser (57) based on 100-way Multiz alignment including Chimp (panTro4), Gorilla (gorGor3), Orangutan (ponAbe2), Gibbon (nomLeu3), Rhesus (rheMac3), Crab-eating macaque (macFas5), Baboon (papHam1), Green monkey (chlSab1), Marmoset (calJac3), Squirrel monkey (saiBol1), Bushbaby (otoGar3), Mouse (mm10), Rat (rn5), Pig (susScr3), Dog (canFam3), Opossum (monDom5), Platypus (ornAna1), Chicken (galGal4), X. tropicalis (Western clawed frog) (xenTro7), Fugu (fr3), Zebrafish (danRer7).

### Data processing and statistical analysis

Data processing was performed using handmade scripts in R (R Foundation for Statistical Computing, Vienna, Austria, https://www.R-project.org/). An R package of qqman (58) was used to make plots. The general threshold of statistical significance was set to 0.05. Used scripts are available upon request.

## RESULTS

### Summary of the analyzed data

We explored the sex-associated DNA methylation in peripheral blood cells using iMETHYL database (**Figure 1A**). The database contains WGBS data from three blood cell types; CD3+/CD4+ T cells (CD4T), CD14++/CD16-monocytes, and CD16+ neutrophils from approximately 50 male and 50 female subjects (30,32). We utilized the mean DNA methylation level and SD for each CpG site on autosomes, calculated separately for males and females within each of the three cell types (**Figure 1B**). Approximately 23 million CpG were used for the analysis after QC, including the removal of approximately 140,000 CpGs that overlapped with common SNPs (**Figure 1C**). Mean methylation levels of CpGs were comparable, ranging from 79.2% to 80.9% across cell types and between sexes (**Figure 1C**).

**Figure 1.**
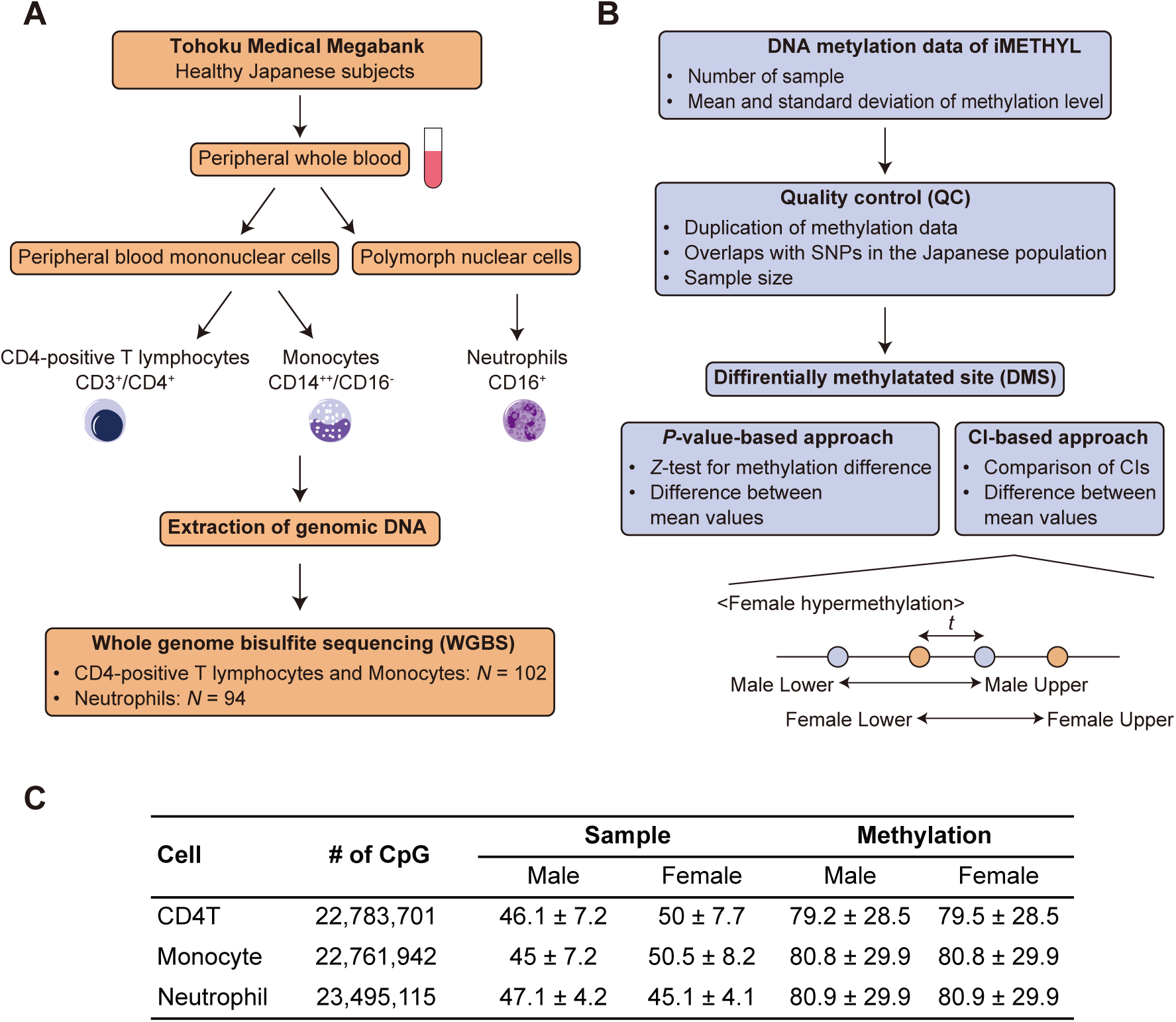
iMETHYL database and strategy for identifying sex-associated DMSs. (**A**) WGBS data in the iMETHYL database. (**B**) Strategy for identifying DMSs. We utilized the mean and standard deviation of DNA methylation levels at each autosomal CpG site, calculated separately for males and females in the three blood cell types. Two approaches were developed for DMS identification. The *P*-value-based approach used *Z*-tests for differences in population proportions with Bonferroni correction. The confidence interval (CI)-based approach compared the 95% CIs of methylation levels between males and females, using *t* to allow a defined degree of overlap between them. In both methods, CpGs showing absolute methylation differences greater than 5% were selected. The illustrated example defines female hypermethylation in the CI-based approach. Male Lower and Male Upper indicate the lower and upper bounds of the 95% CI in males, respectively; Female Lower and Female Upper represent the corresponding bounds in females. (**C**) Summary of WGBS data. # of CpG indicates those remaining after quality control. Mean and standard deviation are shown for sample size and methylation level (%) of each CpG site.

### Identification of DMSs using *P*-value-based approach

We identified a total of 8,013, 4,017, and 5,832 autosomal DMSs for CD4T, monocytes, and neutrophils, respectively, by *P*-value-based approach using *Z*-tests for differences in population proportions with Bonferroni correction (**Figure 2A** and **Supplementary Table S1**). The total number of unique DMSs across cell types was 15,611. Most DMSs were confined to a single cell type, with 82.5%, 71.3%, and 76.6% being specific to CD4T, monocytes, and neutrophils, respectively. The number of hypermethylated DMSs in males (male DMSs) and females (female DMSs) was roughly equal in CD4T (*N* = 4,287 for males and 3,726 for females; male:female ratio = 0.87) and in monocytes (*N* = 2,101 and 1,916; ratio = 0.91), but was high in females compared to males in neutrophils (*N* = 2,170 and 3,662; ratio = 1.69). Both male and female DMSs in three cell types were distributed evenly across the entire autosomes (**Figure 2B**). The mean methylation differences of total male and female DMSs in three cell types were 12.3% and 11.9%, respectively (**Figure 2C**). The distributions of methylation differences were similar across the three cell types and between male and female DMSs (**Figure 2D**).

**Figure 2.**
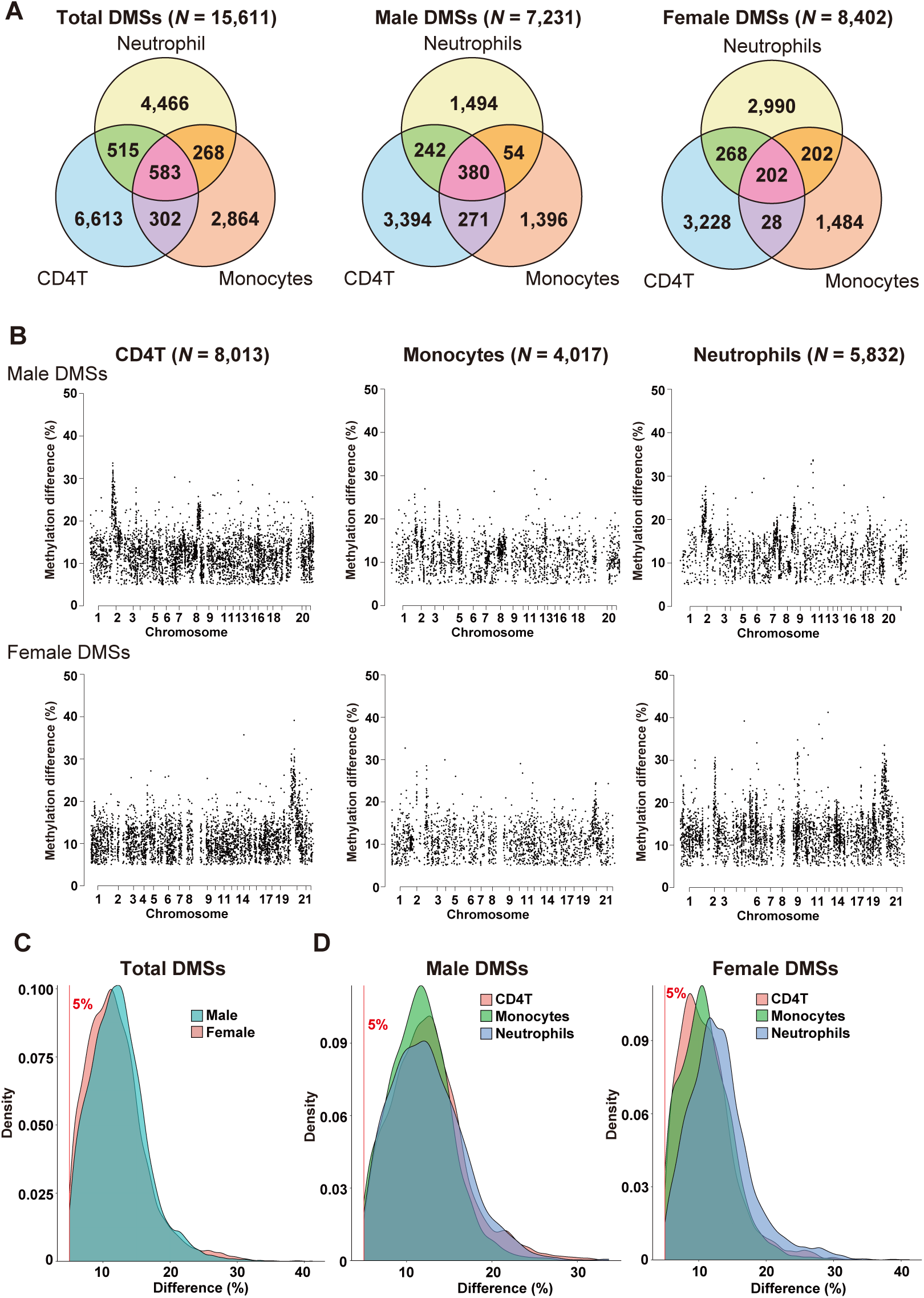
Sex-associated DMSs identified using a *P*-value-based approach. (**A**) Venn diagrams showing overlaps of DMSs among cell types. Total DMSs, along with hypermethylated DMSs in males (male DMSs) and females (female DMSs) are shown. (**B**) Chromosomal distribution of DMSs. Male and female DMSs in CD4T, monocytes, and neutrophils are shown. (**C**, **D**) Density plots of methylation differences. Total DMSs (C), male and female DMSs (D) are shown. For total DMSs, methylation differences in male and female DMSs are displayed separately. For male and female DMSs, differences in the three cell types are also shown separately. The 5% line indicates the threshold used in the analysis.

### Identification of DMRs

We defined DMRs by merging DMSs located within 1 kb of each other, and identified a total of 1,657 DMRs (**Table 1** and **Supplementary Table S2**). The number of DMRs ranged from 168 to 444 in three cell types. The median lengths were between 134 and 217 bp, with longer DMRs in neutrophils.

**Table 1.**
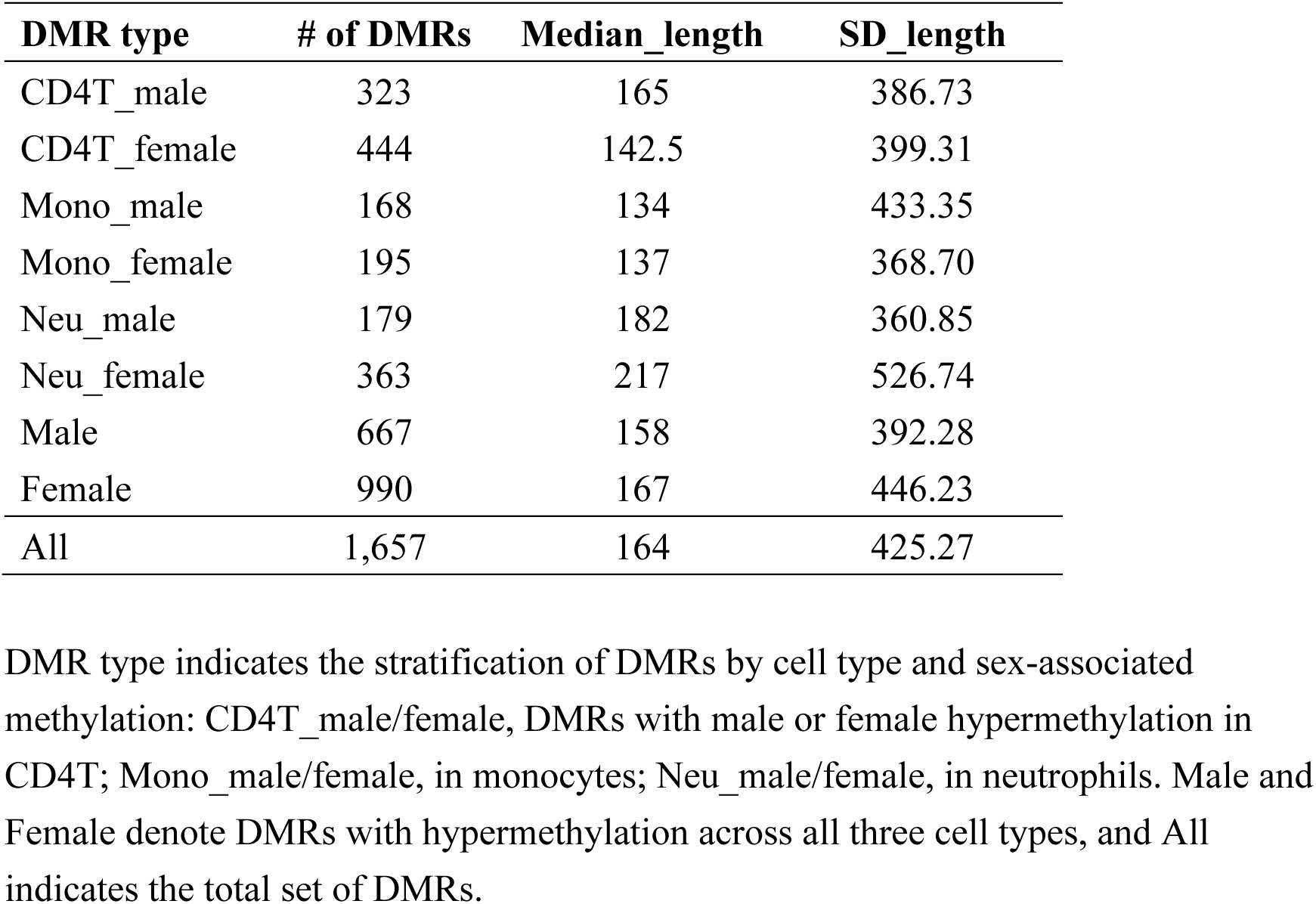
Summary of DMRs identified by *P*-value-based approach.

### Identification of DMSs using CI-based approach

To identify robust DMSs, we introduced CI-based approach using the conventional Wald method, and obtained a total of 126 DMSs from three cell types. We then introduced a parameter *t*, which represents the tolerance for the overlap between 95% CIs of males and females (**Figure 1B** and Materials and Methods). As expected, the number of CpGs identified as sex-associated DMSs increased with higher *t* values (**Figure 3A**), while the mean DNA methylation differences decreased (**Figure 3B**). We then evaluated the effect of *t* by calculating the average methylation differences in CpGs selected using a sliding window approach. To determine the optimal value of *t*, we searched the point of the steepest gradient and set *t* to 2.2% (**Figure 3C**). Finally, we identified a total of 120, 6, and 128 autosomal DMSs for CD4T, monocytes, and neutrophils, respectively (**Figure 3D**). The number of unique DMSs was 213. The majority of DMSs were cell-type specific and distributed across the entire autosomes (**Figure 3D, E** and **Supplementary Table S3**). As expected, most of which (96.7%) were included in the *P*-value based approach. The increased number of female DMSs in neutrophils (*N* = 22 for male and 106 for female; ratio = 4.88) was also observed. The mean differences of total male and female DMSs in three cell types were 25.7% and 27.3%, respectively (**Figure 3F**). We also identified 22 DMRs in CI-based approach (**Supplementary Table S4**).

**Figure 3.**
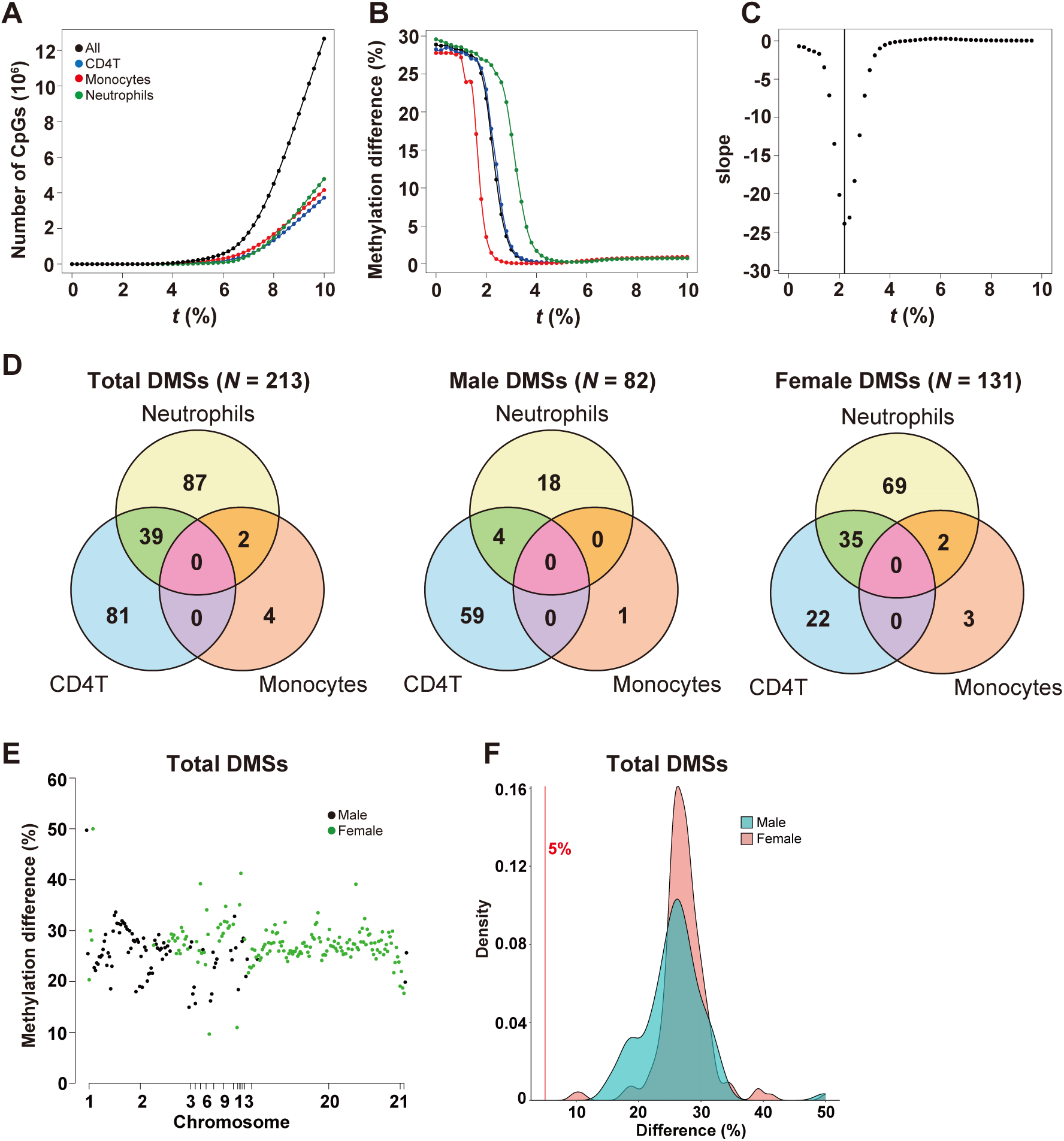
Establishment of the CI-based approach and DMS distribution (**A**) Relationship between *t* and the number of DMSs. *t* specifies the permitted overlap between 95% CIs of male and female methylation levels. (**B**) Relationship between *t* and average methylation difference. (**C**) Methylation differences calculated using a sliding window of the five nearest plots in three cell types. The steepest gradient was used to determine the optimal *t*, which was set to 2.2%. (**D**) Venn diagrams showing overlaps of DMSs among cell types. (**E**) Chromosomal distribution of DMSs identified by CI-based approach. The horizontal and vertical axes represent the chromosomal location and the methylation difference (%) of each DMS with male (black) or female (green) hypermethylation, respectively. (**F**) Density plots of methylation differences. For total DMSs, methylation differences in male and female DMS are displayed separately. The 5% line indicates the threshold used in the analysis.

### Genic and CpG island annotation of DMSs

We characterized the genomic distribution of the DMSs by *P*-value-based approach by annotating their locations with respect to gene structure and CpG island context (**Figure 4A**). The majority of DMSs were located in intronic regions (35.5%), and when including exonic regions (11.1%) and UTRs (3.4%), DMSs within gene bodies accounted for 50% of the total. The next most frequent location was regulatory regions, including promoters and 1-5 kb upstream from the transcription start site (TSS), which comprised 20.2%. Notably, intergenic regions contained relatively a small proportion of DMSs (29.9%). In terms of CpG islands, DMSs were almost evenly distributed between CpG islands including their flanking regions, and non-CpG island regions (inter CpG island). These proportions were broadly consistent regardless of cell type or sex-specific classification. When CI-based DMSs were used for the analysis, enrichment in intronic regions became more prominent (41.3%), whereas enrichment within CpG islands and their flanking regions was markedly diminished (18.3%) (**Figure 4B**). GO analysis of the DMS-associated genes revealed significant enrichment of synaptic and cell adhesion functions, irrespective of cell type or sex (**Figure 4C** and **Supplementary Table S5**). Notably, blood function-related GO terms were specifically enriched in female DMSs in CD4T, such as “T cell activation” (adjusted *P*-value = 9.6 × 10^-12^) and “T cell differentiation” (adjusted *P*-value = 1.3 × 10^-8^). GO analysis with correction for gene length confirmed enrichment of neuronal functions in DMS-associated genes other than female CD4T, while blood function-related terms were specifically enriched in female CD4T (**Supplementary Table S6**).

**Figure 4.**
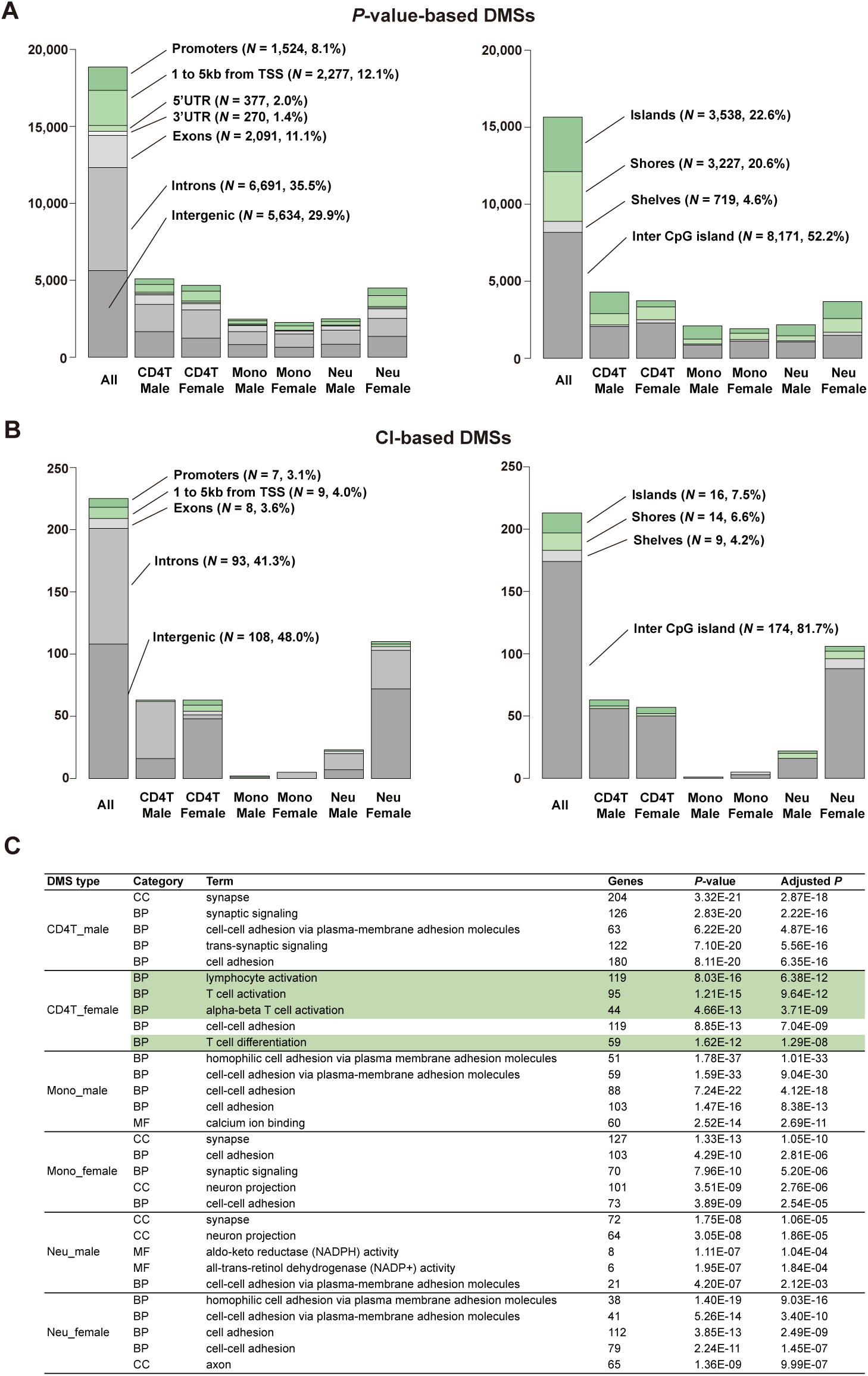
Annotation of DMSs and their associated genes. (**A**) Annotation of DMSs identified by *P*-value-based approach in relation to genic (left) and CpG island (right) context. (**B**) Annotation of DMSs identified by CI-based approach of genic (left) and CpG island (right) context. (**C**) GO analysis of the DMS-associated genes by *P*-value-based approach. The top 5 terms are shown. The full list is available in **Supplementary Table S5**, and the results corrected for gene length are shown in **Supplementary Table S6**. GO terms associated with blood functions are highlighted in green. Genes indicate the number of the DMS-associated genes. Adjusted *P* indicates *P*-value with Bonferroni correction. TSS, transcription start site. CC, cellular component; MF, molecular function; BP, biological process.

### Effects of imprinting and smoking on sex-associated methylation

Imprinting is known to influence DNA methylation on autosomes. We evaluated the overlap between the DMSs identified by *P*-value-based approach and reported imprinting control regions (ICRs) (59). Only 31 DMSs overlapped with ICRs, indicating that the observed methylation differences were not substantially influenced by imprinting. We also examined the overlap with smoking-associated CpG sites. Among the 972 smoking-related CpGs reported previously (38), only 12 overlapped with our DMSs. These findings suggest that smoking had little impact on the sex-associated methylation identified in this study.

### Comparison with previously reported sex-associated DMSs in blood

We compared sex-associated DMSs in eight previous studies (19,20,23–28) with those identified by *P*-value-based approach in the present study (**Figure 5A**). Nearly all DMSs reported in the eight studies were covered by our WGBS analysis (97.3-100%). We found that only a small fraction of DMSs was validated, with validation rates ranging from 0.1% to 8.7% at the DMS level and from 0.2% to 13.0% at the DMR level in six studies. Two studies (23,24) showed relatively higher validation rates, with 24.8% and 55.6% at the DMS level, and 35.0% and 62.2% at the DMR level, respectively. We plotted the methylation differences of previously reported DMSs using our dataset (**Figure 5B, C**). In the six studies with low validation rates, most of the reported DMSs showed methylation differences of less than 5% in our dataset (**Figure 5B**). By contrast, the two studies with high validation rates showed relatively larger methylation differences and a greater density of DMSs in neutrophils (**Figure 5C**).

**Figure 5.**
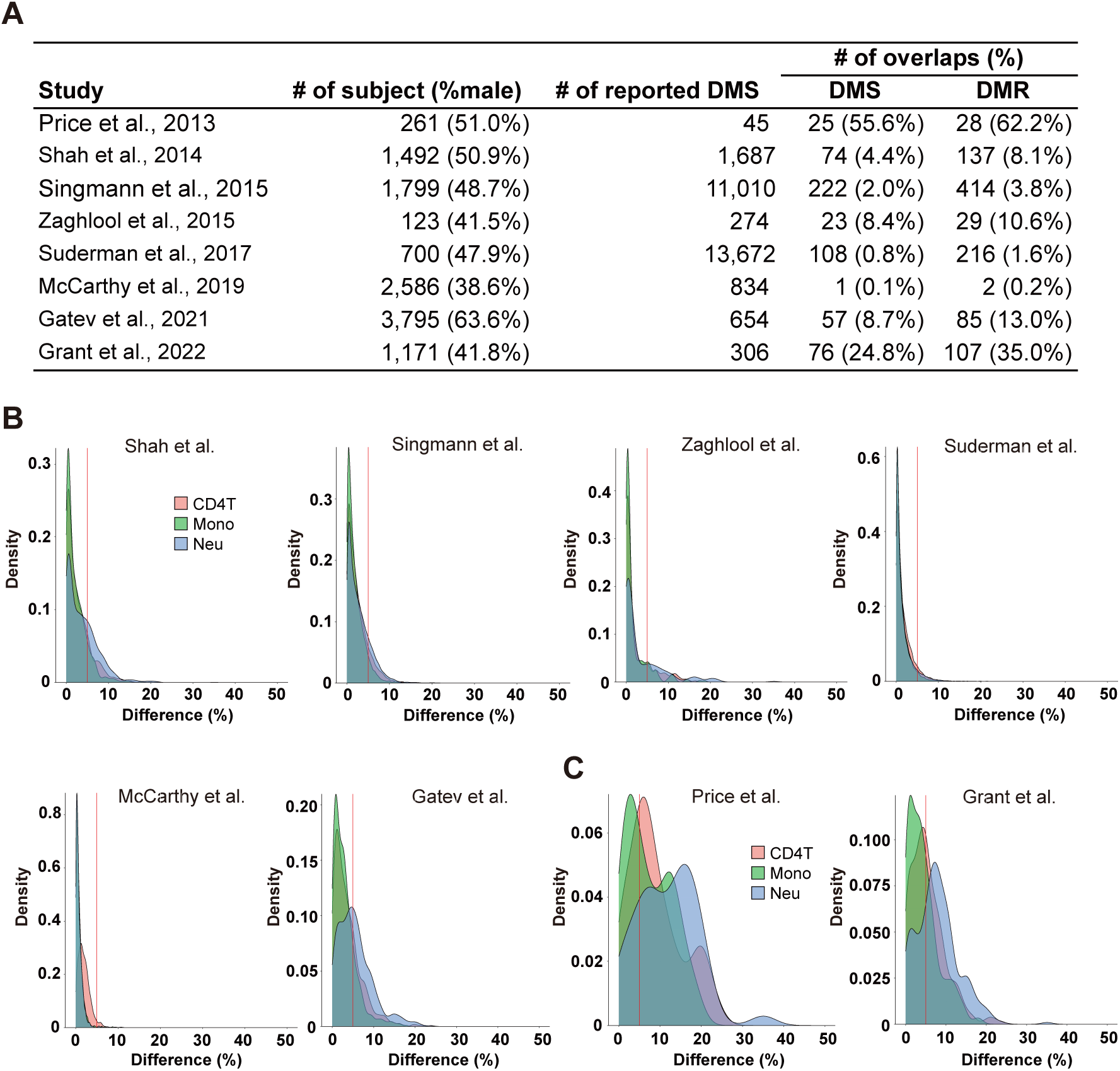
Methylation differences of previously reported DMSs in our dataset. (**A**) Comparison of sex-associated DMSs with previous studies. (**B**, **C**) Density plots show methylation differences of previously reported sex-associated CpG sites, with a reference line (red) at 5%. The datasets for (B) were derived from studies with low levels of replication (20,22,25–28), whereas those for (C) came from studies with high levels of replication (23,24). The horizontal axis indicates the methylation difference (%) between males and females in our dataset.

### Enrichment in reported GWAS regions

We collected locus-level GWAS data for various traits from the GWAS Catalog and examined whether DMRs by *P*-value-based approach were significantly enriched within these loci. We tested a total of 337 GWASs selected from approximately 130,000 entries. Nominally significant enrichment with one of the six types of DMRs was observed for a total of 75 GWAS traits, of which 65 were unique, and three of them remained significant after correction for multiple testing (**Table 2** and **Supplementary Table S7)**. Female DMRs in neutrophil were significantly enriched in GWAS regions of lifetime smoking (60) (OR = 35.4, adjusted *P*-value = 0.013) and psoriasis (61) (OR = 10.1, adjusted *P*-value = 0.025). For lifetime smoking, four DMRs, including two genes (*SLC4A10* and *RASGRF2*), overlapped with GWAS regions. For psoriasis, six DMRs including four genes (*RGS14*, *TOB2P1*, *AGPAT1*, and *TSBP1*) overlapped with GWAS regions. In addition, female DMRs in CD4T were significantly enriched in GWAS regions of multiple sclerosis (MS) (62) (OR = 8.8, adjusted *P*-value = 0.020), with seven DMRs including three genes (*JADE2*, *RGS14*, and *DLEU1*).

**Table 2.**
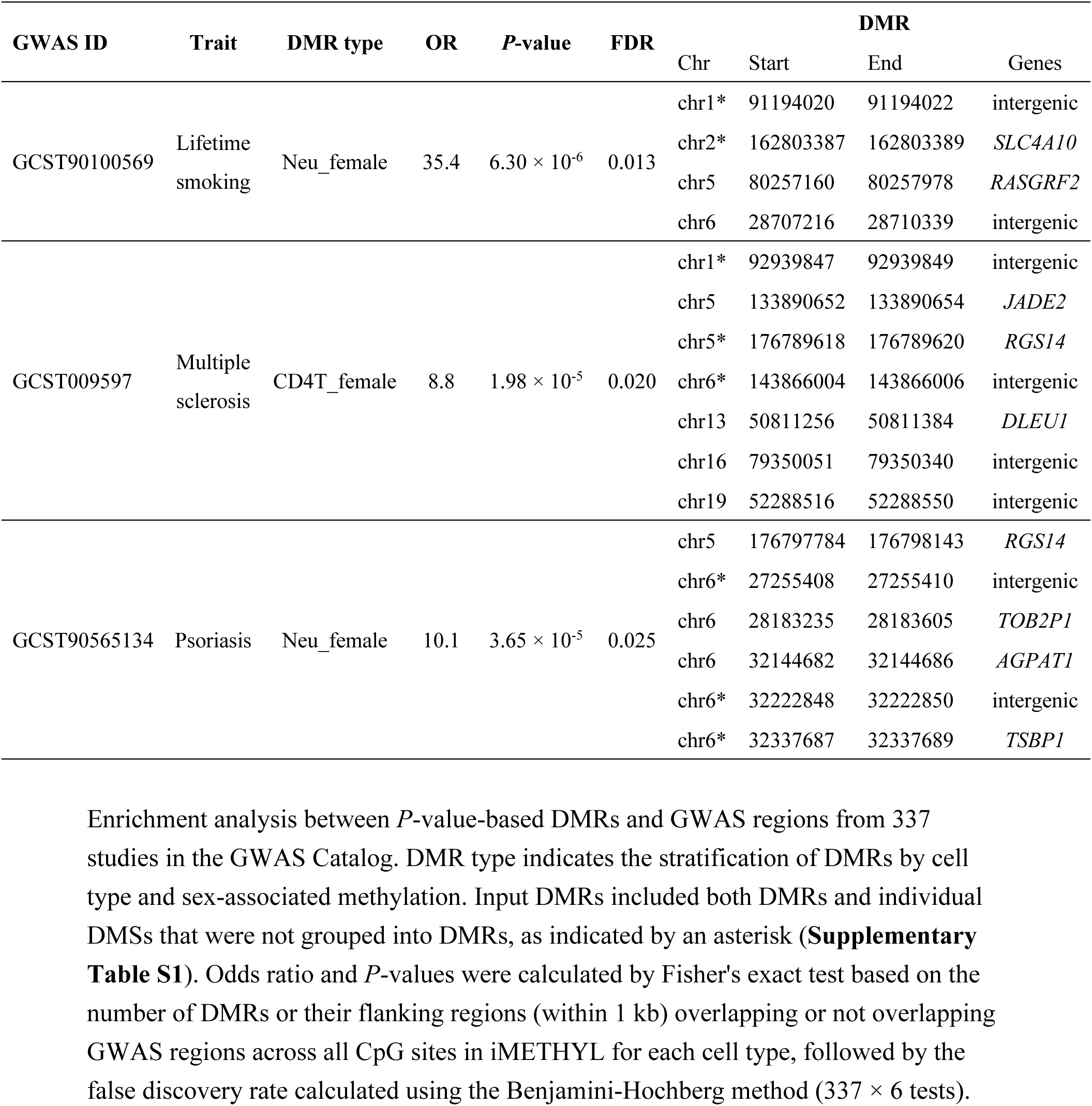
Enrichment of DMRs in reported GWAS regions.

### Example of expression level of DMS-associated gene

We further focused on the relationship between gene expression level and methylation status. As a representative example, *RIMBP3*, one of the genes annotated with CI-based DMSs, showed male hypermethylation associated with lower expression in male whole blood in the GTEx database (51) (Mann-Whitney *U* test, *P*-value = 0.029) (**Figure 6A, B**). *RIMBP3* is a single-exon gene with a CpG island at its 5′ end. Significant male hypermethylation was observed in the CpG island in neutrophils, with a similar trend in other cell types, whereas no sex differences were detected in other regions (**Figure 6A** and **Supplementary Figure S1**).

**Figure 6.**
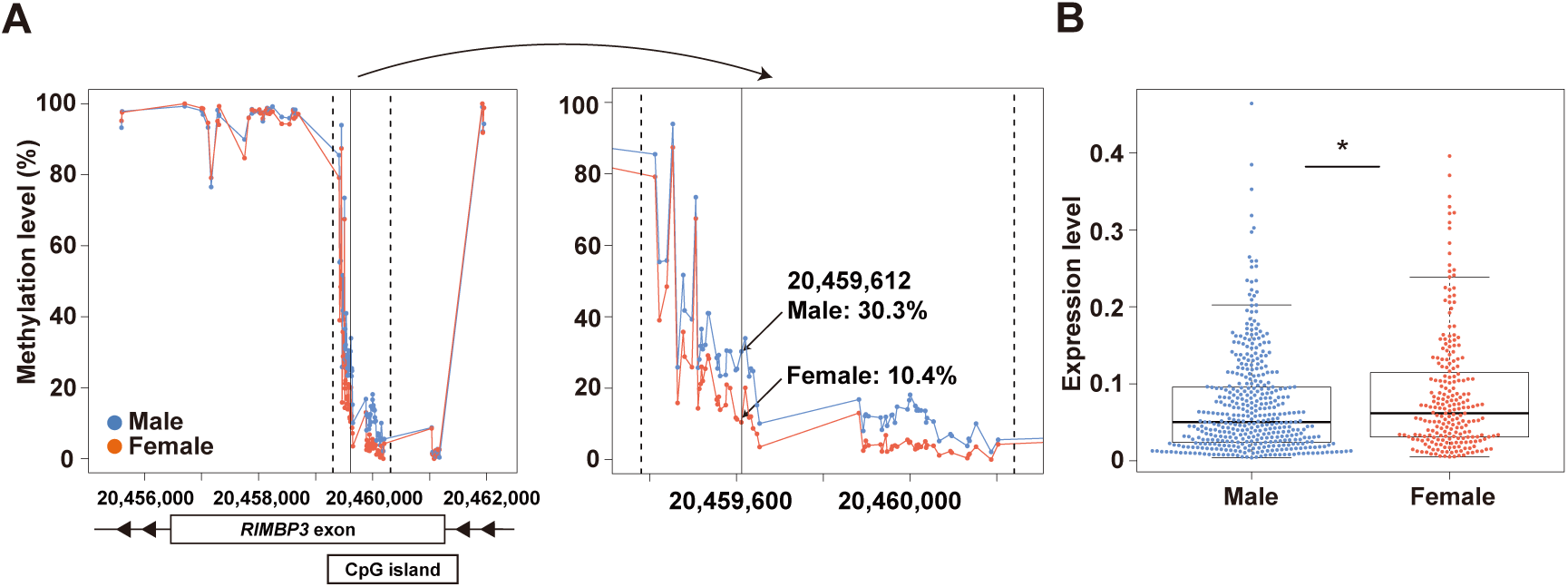
*RIMBP3* expression and methylation profiles. (**A**) Methylation levels of CpG sites in both the exon of *RIMBP3* and CpG island are shown on the left, including a CpG site exhibiting male hypermethylation in neutrophils (hg19 coordinates, chr22:20,459,612) with a solid line. The detailed methylation status within the dashed lines is shown on the right. (**B**) Expression level of *RIMBP3* based on public RNA-seq data in whole blood from 424 males and 227 females. The asterisk indicates statistical significance (*P*-value < 0.05).

### TF target sequences

To evaluate the genome-wide relationship between DMSs and TF target sequences, we conducted enrichment analysis using ChIP-Atlas, which incorporates previously published ChIP-seq data (52–54). Among the six DMS types, we detected significant enrichment of TFs in the male and female DMSs in CD4T and neutrophils (**Figure 7** and **Supplementary Table S8**). The TFs common to all four DMS types were EGR1, ELF1, ERG, KDM2B, KMT2A, RUNX1, and TRIM28. They were either chromatin regulators (KDM2B, KMT2A, TRIM28) or hematopoietic-related TFs (EGR1, ELF1, ERG, RUNX1), reflecting a core regulatory network established early in hematopoiesis. In female DMSs in CD4T, we found enrichment of regulators of T cell differentiation and development, such as MYB and TCF7, which was consistent with the unique enrichment of GO terms for DMS-associated genes. In addition, PPI analysis revealed that the commonly enriched TFs acted as network hubs, around which other TFs with related functions in hematopoiesis, blood cell regulation, and immune functions were organized, suggesting the presence of coordinated regulatory networks (**Figure 8** and Supplementary Table S9).

**Figure 7.**
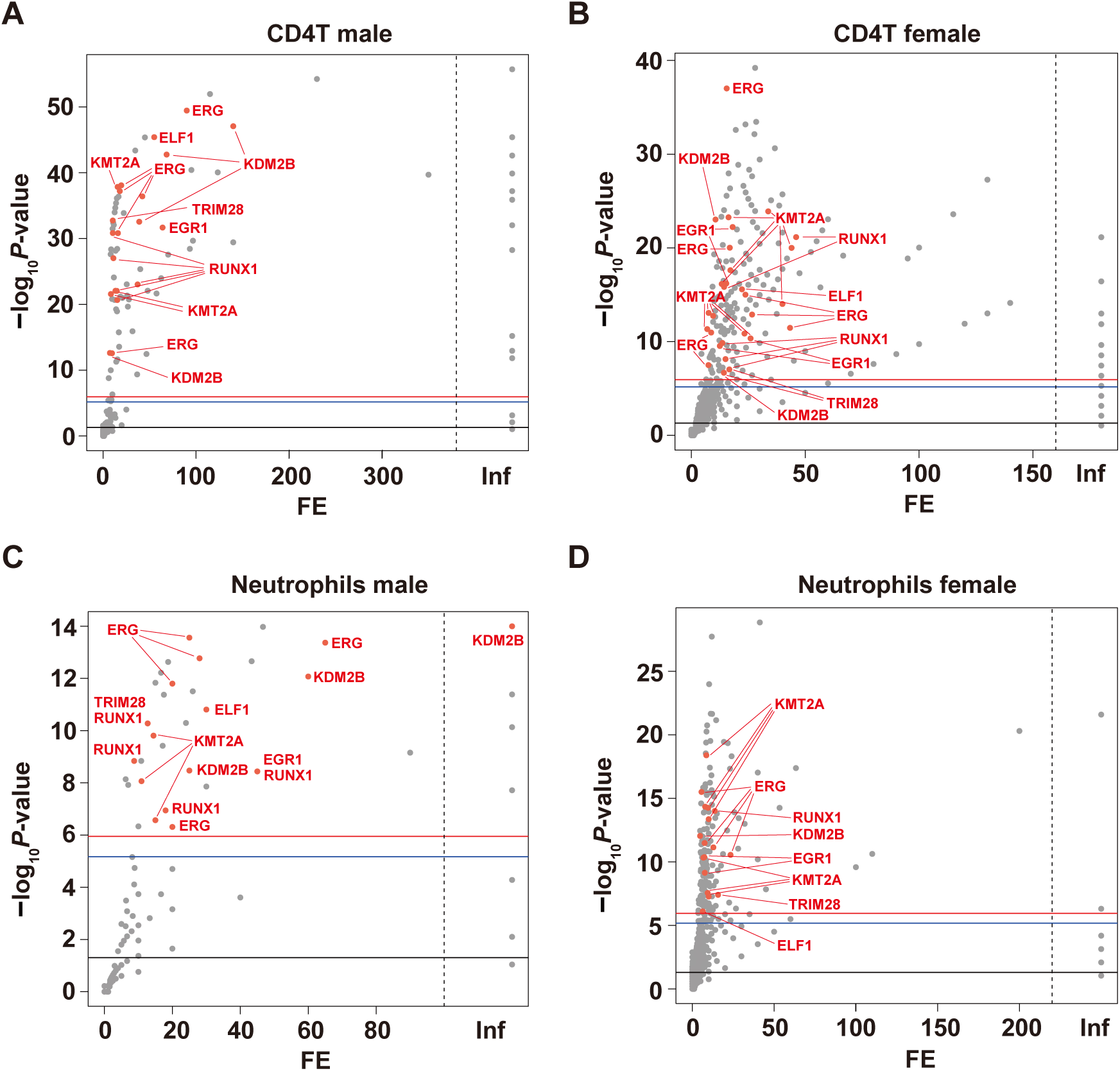
Enrichments of DMSs in transcription factor target sequences. Volcano plots showing the enrichment of TF target sequences around DMSs identified by CI-based approach in each DMS type. Analyses were performed for DMSs in CD4T male (**A**) CD4T female (**B**), neutrophils male (**C**), and neutrophils female (**D**). Full list is available in **Supplementary Table S8**. The horizontal axis indicates fold enrichment. The red plots indicate common TFs between all four DMS types. ‘Inf’ denotes cases where enrichment analysis of certain TFs showed no overlaps with the background data. The black, blue, and red lines represent significance thresholds after multiple testing using Fisher’s exact test: uncorrected *P*-value < 0.05 (black), Bonferroni correction for 7,397 tests (blue), and Bonferroni correction for 7,397 × 6 DMS types (red).

**Figure 8.**
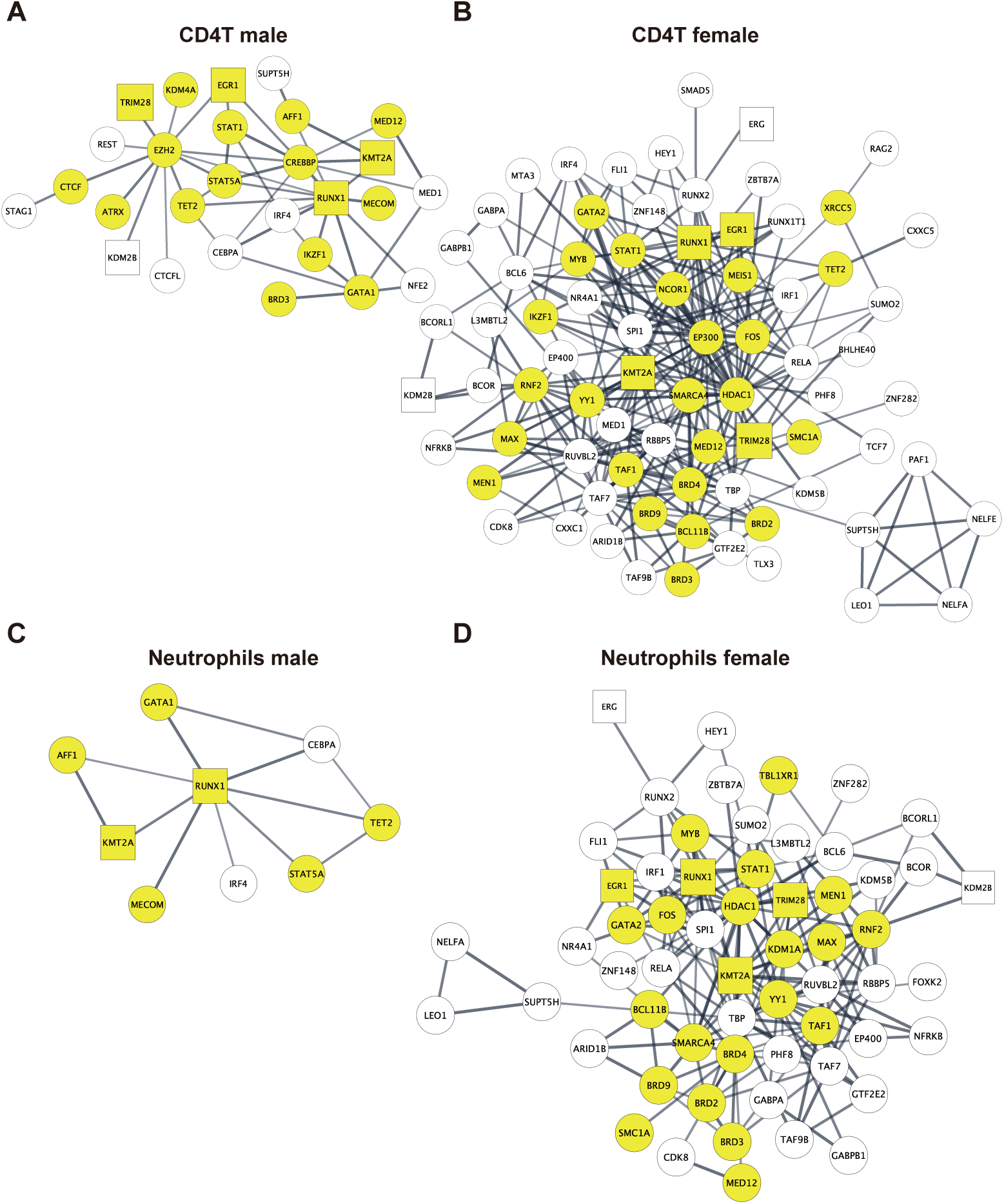
PPI analysis of TFs associated with DMSs. PPI analyses were performed using TFs enriched in DMSs in CD4T male (**A**), CD4T female (**B**), neutrophils male (**C**), and neutrophils female (**D**). Detailed list is available in **Supplementary Table S9**. Hub nodes related to leukemia cell are highlighted in yellow. The square nodes indicate common TFs between all four DMS types.

### Conserved CpG sequences of the DMSs

We examined whether sequences of sex-associated CpGs are conserved through evolution. Among the 213 DMSs identified by CI-based approach, we observed that the number of conserved sequences increased in species closely related to humans but decreased markedly in rodents, and little conservation was observed in marsupials, monotremes, birds, amphibians, or fish (**Figure 9A**). Notably, one site was fully conserved, including surrounding sequences, across mammals such as primates and rodents, and conservation extended to the CpG site itself in all species except fish. This site is in the second intron of *FIGN* (**Figure 9A**), and showed female hypermethylation in neutrophils (**Figure 9B**), with a consistent change in monocytes but not in CD4T (**Supplementary Figure S1**). We investigated *FIGN* expression levels using whole blood datasets and observed lower expression in females, consistent with female hypermethylation (Mann-Whitney *U* test, *P*-value < 0.01) (**Figure 9C**).

**Figure 9.**
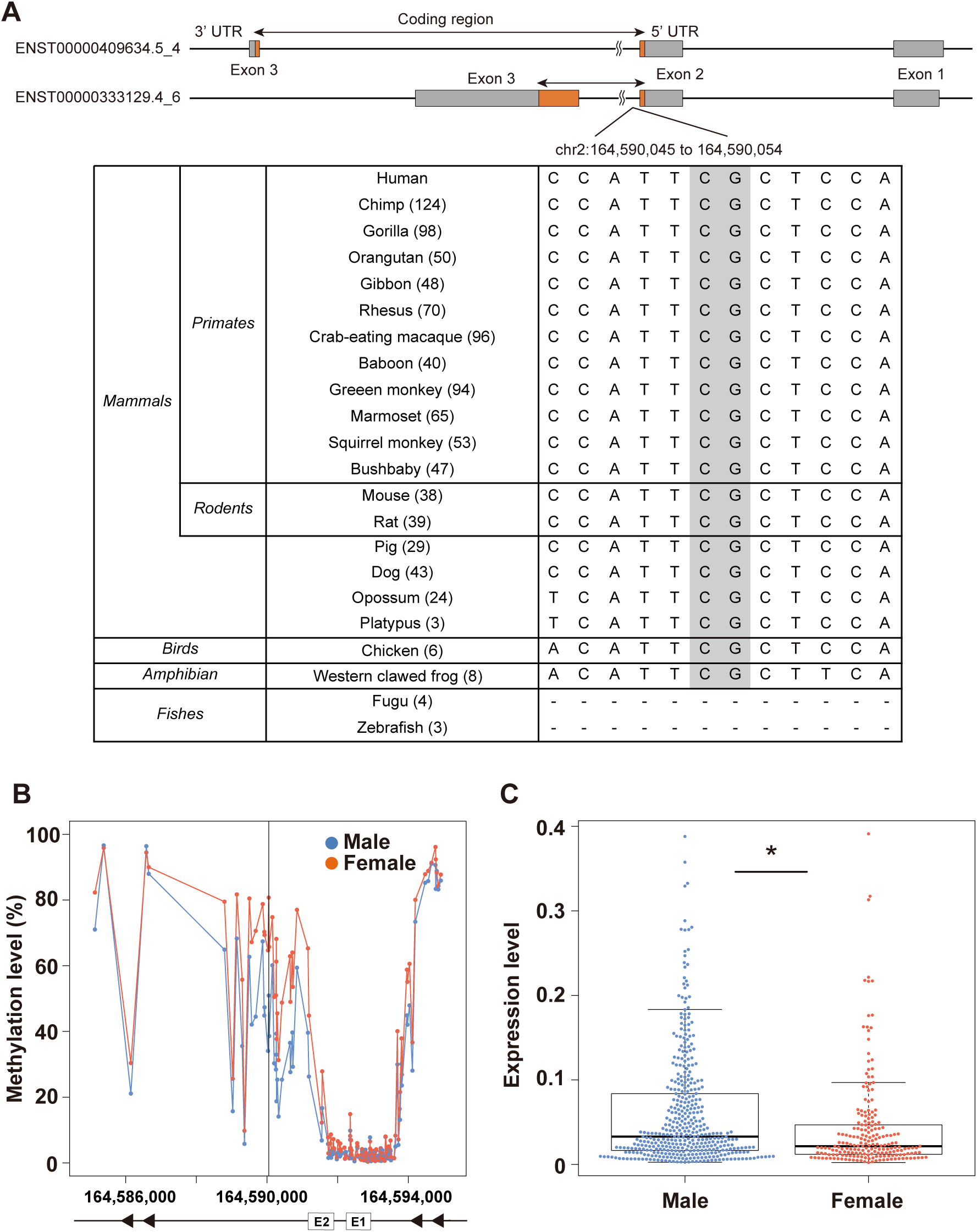
Conserved CpG sequence of DMS in the *FIGN* gene across evolution, and its methylation and expression profiles. (**A**) Transcripts of human *FIGN* on the minus strand are shown, with the coding regions colored in brown. One CpG site, located in the second intron (hg19 coordinates, chr2:164,590,051), is conserved. The notation ‘-’ denotes missing or unalienable sequences in Fugu and Zebrafish. The number of conserved CpG sequences in each species is shown in parentheses. (**B**) Methylation status around a conserved CpG site in *FIGN* in neutrophils. A black line indicates the conserved CpG site. E1 and E2 indicate the first and second exon of *FIGN*, respectively. (**C**) Expression levels of *FIGN* based on public RNA-seq data in whole blood from 417 males and 207 females. The asterisk indicates statistical significance (*P*-value < 0.05).

## DISCUSSION

This study established the first genome-wide, cell type-resolved map of sex-associated DNA methylation in peripheral blood. By applying WGBS with two complementary analytical approaches, we overcame limitations of prior array-based studies and revealed thousands of autosomal sites with robust sex differences. These differences were largely cell type-specific, with female-biased hypermethylation in neutrophils. Functional annotation highlighted gene body enrichment, implicating neuronal and immune pathways, while TF binding site analysis indicated that such differences may be established early during hematopoietic differentiation. Importantly, DMRs were significantly enriched in GWAS loci for smoking, psoriasis, and MS, supporting their potential relevance to disease susceptibility. Evolutionary conservation analyses revealed broadly conserved CpGs, such as in *FIGN*, and lineage-restricted conservations across vertebrates. In addition, an example of *RIMBP3* demonstrated concordant methylation and expression differences between sexes. Collectively, these findings expand our understanding of autosomal epigenetic regulation in shaping sex differences and provide a foundation for advancing sex-informed biology and medicine.

### Cell type-specific features and sex differences in methylation

Our analysis revealed that the majority of sex-associated DMSs were unique to a single cell type, underscoring the importance of cell-type resolution in epigenetic studies. Previous studies using whole blood suggested largely female-biased hypermethylation (19,20,23). In contrast, our results demonstrate that this pattern is not uniform across cell types. CD4T cells and monocytes showed relatively balanced numbers of male and female DMSs, whereas neutrophils exhibited a pronounced female bias in hypermethylated DMSs, suggesting a stronger epigenetic sexual dimorphism in this cell population.

### Enrichment of neuronal functions in DMS-associated genes

An unexpected result was the enrichment of neuronal and synaptic functions among genes annotated to sex-associated DMSs. Because our study was conducted in blood cells and the overall methylation patterns were strongly cell type-specific, this enrichment is unlikely to indicate a direct similarity between blood and brain methylomes. Instead, it is more likely that many genes categorized as neuronal have pleiotropic roles, such as adhesion, cytoskeletal remodeling, and intracellular signaling, that are also essential for hematopoietic cells. This interpretation is consistent with our TF-binding site analysis, which indicated enrichment of TFs related to hematopoiesis and immune regulation rather than neural development. At the same time, the recurrence of neuronal gene categories is noteworthy given that several of these loci have also been implicated in GWAS of traits and disorders closely related to neural function and neurodevelopment. Although causality cannot be inferred, such findings raise the possibility that peripheral methylation signatures contribute to understanding sex differences in susceptibility to brain-related traits.

### Comparison with the previous studies

When compared with eight previous array-based studies of sex-associated DNA methylation in blood, nearly all of the reported CpGs were covered by our WGBS dataset. However, only a small fraction of these sites met our criteria for sex-associated differences, with validation rates generally below 10%. Two prior studies achieved higher replication, suggesting that certain analytic thresholds and cell-type contributions can strongly influence reproducibility. The limited overlap indicates two major considerations. First, differences in analytic thresholds, combined with the restricted genomic coverage of array-based platforms, can substantially affect the detection of sex-associated CpGs. In fact, only about 7% of the CpGs identified as DMSs are covered by the EPICv1 array (**Supplementary Figure S2**). Second, the use of whole blood in previous work introduces confounding from cellular composition, leading to apparent discrepancies when signals are cell type-restricted. Our cell type-resolved analysis reveals that many sex-associated differences are diluted or missed entirely in mixed-cell datasets.

### Convergence of epigenetic sex differences and GWAS loci

The enrichment of sex-associated DMRs within GWAS loci suggested the potential functional relevance of blood cell methylation differences in traits and diseases with known sex biases. Although these associations cannot establish causality, they suggest that sex-associated epigenetic regulation may converge with genetic susceptibility to influence phenotypic outcomes.

Among the enriched GWAS loci, lifetime smoking showed the strongest association with female-hypermethylated DMRs in neutrophils. Smoking behavior exhibits well-documented sex differences in prevalence, initiation age, and cessation rates, and sex hormones have been implicated in modulating nicotine metabolism and dependence (63–65). Overlapping genes included *SLC4A10*, which encodes a sodium bicarbonate transporter expressed in the brain and implicated in neuronal excitability (66) and *RASGRF2*, a guanine nucleotide exchange factor involved in synaptic plasticity and learning (67). The neutrophil-specific enrichment may reflect not only genetic predisposition but also downstream consequences of smoking exposure, such as sex-dependent differences in inflammatory responses.

We also observed significant enrichment of female-hypermethylated DMRs in CD4T cells within GWAS loci associated with MS. MS is a prototypical autoimmune disorder with a strong female predominance, and both genetic susceptibility and immune dysregulation contribute to its pathogenesis (68). Overlapping genes included *JADE2*, *RGS14*, and *DLEU1*. *JADE2* encodes a component of a histone acetyltransferase complex (69) that may influence T-cell transcription through chromatin remodeling. RGS14 modulates G protein and MAPK signaling, thereby contributing to lymphocyte signaling dynamics (70). *DLEU1* is a long non-coding RNA gene that regulates NF-κB activity in immune cells, and its transcription activity is regulated by DNA methylation in the promoter region (71). Female-biased methylation within these loci suggests that epigenetic differences in T cells could influence immune pathways underlying MS susceptibility.

We also identified enrichment of female-hypermethylated DMRs in neutrophils within GWAS loci for psoriasis. Psoriasis is a chronic inflammatory skin disorder with modest female predominance in some populations (72,73). Overlapping genes included *RGS14*, *TOB2P1*, *AGPAT1*, and *TSBP1*. *RGS14*, as discussed above, has been implicated in lymphocyte signaling. *TOB2P1* belongs to the TOB/BTG family of antiproliferative proteins that showed altered expression in inflammatory bowel disease (74). *AGPAT1* encodes an enzyme involved in phospholipid biosynthesis that may affect membrane signaling and inflammatory mediator production (75). *TSBP1* is a poorly characterized gene located within the MHC class II region, where many immune-related susceptibility loci reside. The neutrophil specificity of this enrichment suggests that sex-associated methylation could modulate inflammatory responses relevant to psoriasis pathogenesis.

### TF-binding site enrichment suggests early establishment of sex-associated methylation

TF-binding site analysis indicates that sex-associated methylation is embedded within transcriptional networks through regulatory regions bound by TFs that govern lineage identity and chromatin state. This implies that such epigenetic differences are established early in blood cell differentiation and maintained within immune regulatory networks. Network-level analyses further implied that enriched TFs serve as hubs connecting chromatin modifiers and immune regulators, raising the possibility that sex-associated methylation contributes to coordinated regulatory circuits rather than single-gene effects.

### Evolutionary conservation of sex-associated CpG sites

We found that only a limited subset of DMSs was conserved beyond primates, with a sharp decline in conservation observed in rodents and virtually no conservation in more distantly related vertebrates. This pattern is consistent with the notion that sex-associated methylation has evolved relatively recently and may contribute to lineage-specific traits in primates. Notably, the CpG site within the intron of *FIGN* was conserved across mammals and even present in birds and amphibians. *FIGN* encodes the protein fidgetin, and is a member of the AAA protein family with functions as a microtubule-severing enzyme in humans, playing roles in cell cycle regulation, cell migration, and neural function (76). A meta-analysis reported sex-specific methylation trajectories with aging at CpG sites within this gene, showing female hypermethylation (77). Female-biased hypermethylation at this gene, accompanied by reduced gene expression in females, suggests a functional importance of epigenetic differences. Overall, these findings indicate that while most sex-associated methylation patterns are evolutionarily labile, a small number of conserved CpGs may play roles in preserving sex-differential regulation across vertebrate lineages.

### Limitations

Several limitations of this study should be acknowledged. First, our analyses relied on summarized statistics provided by the iMETHYL database rather than individual-level sequencing data. This precluded adjustment for inter-individual variability and covariates such as lifestyle or environmental exposures. Second, while three major blood cell types were analyzed, other immune subsets and non-hematopoietic cells were not included, leaving open the possibility that additional sex-associated methylation patterns remain undiscovered. Third, we did not address how sex-associated methylation might arise during development or change across the life course. Fourth, although we observed concordance between methylation and gene expression at selected loci such as *RIMBP3* and *FIGN*, transcriptomic validation was limited to publicly available bulk RNA-seq data from whole blood. Finally, our evolutionary comparisons were based on sequence conservation, and functional conservation of methylation at orthologous sites remains to be experimentally demonstrated.

Despite these limitations, our study provides a comprehensive map of sex-associated DNA methylation across major peripheral blood cell types. Our findings serve as a foundation for future studies exploring the mechanisms and clinical implications of sex-specific epigenetic variation in health and disease.

## ACKNOWLEDGEMENTS

We are deeply grateful to Yayoi Otsuka-Yamasaki for her dedicated efforts in preparing the ethical review application and facilitating the internal collaboration procedures for this study. The Genotype-Tissue Expression (GTEx) Project was supported by the Common Fund of the Office of the Director of the National Institutes of Health, and by NCI, NHGRI, NHLBI, NIDA, NIMH, and NINDS.

## AUTHOR CONTRIBUTIONS

Yutaro Yanagida: Conceptualization, Formal analysis, Data curation, Methodology, Visualization, Validation, Writing original draft, Writing review & editing. Yutaka Nakachi: Conceptualization, Supervision, Project administration, Methodology, Validation, Writing review & editing. Miki Bundo: Supervision, Validation, Writing review & editing. Syohei Komaki: Formal analysis, Resources, Data curation, Validation, Writing review & editing. Atsushi Shimizu: Formal analysis, Resources, Data curation, Validation, Writing review & editing. Kazuya Iwamoto: Conceptualization, Supervision, Project administration, Validation, Writing original draft, Writing review & editing.

## CONFLICT OF INTEREST

None declared.

## FUNDING

This work was partly supported by the Japan Society for the Promotion of Science (JSPS) Grants-in-Aid for Scientific Research (KAKENHI) JP22K07583, JP23K27531, JP25H01314, JP25H01309, JP23H03838, JP24K22098, and JP25K10839, and by Japan Agency for Medical Research and Development (AMED) JP19dm0207074 and JP24wm0625302. The funder had no role in the conceptualization, design, data collection, analysis, decision to publish, or preparation of the manuscript.

## DATA AVAILABILITY

The scripts used in the present study are available from the corresponding author on request.

## Notes

### Competing Interest Statement

The authors have declared no competing interest.

