## Supplementary Figures for "Genome-wide mapping of sex-associated autosomal DNA methylation in major blood cell types"

**Supplementary Figure S1.**

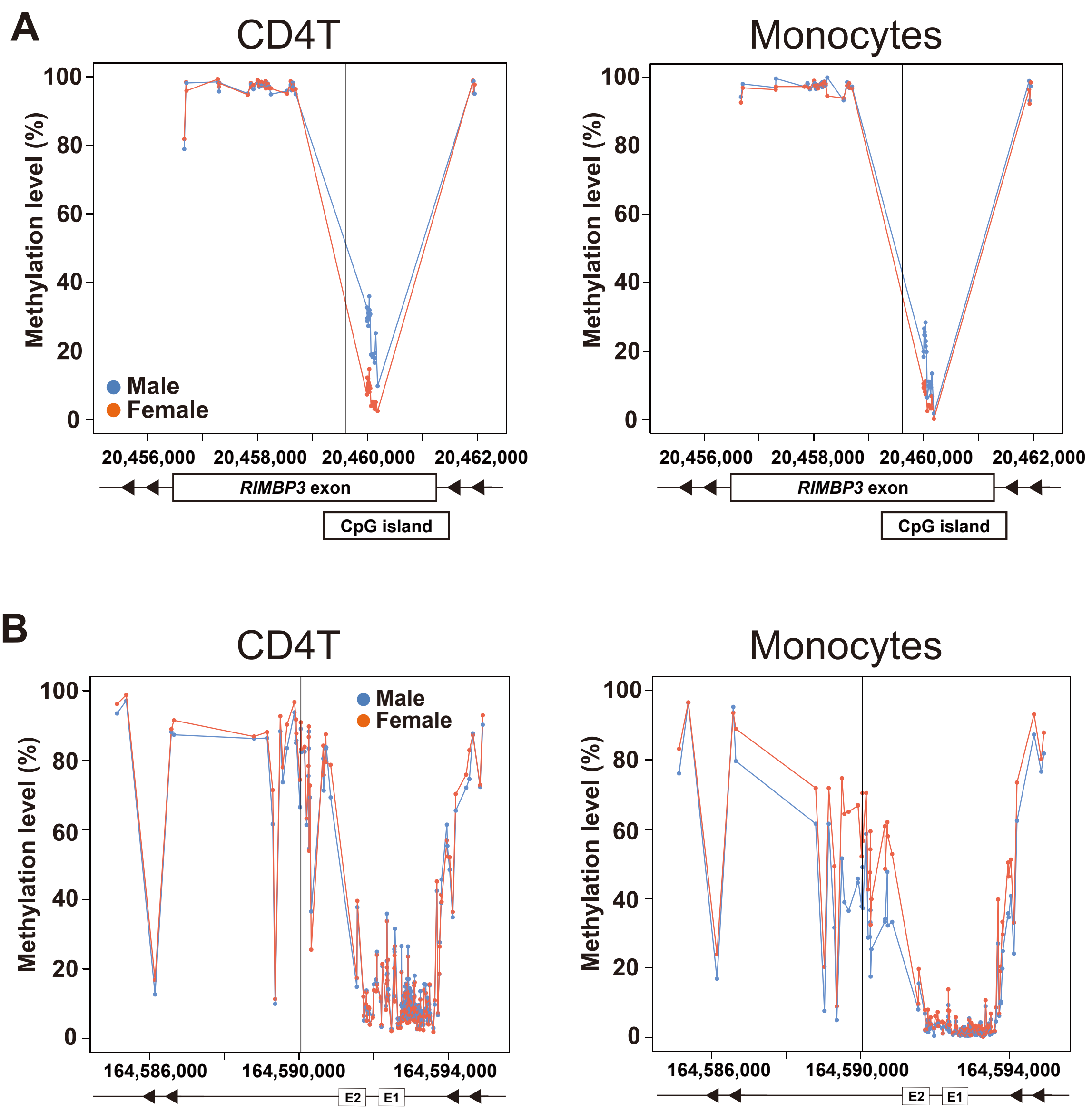


**Supplementary Figure S1.** Methylation status of *RIMBP3* and *FIGN* in CD4T and monocytes. (**A**) Methylation status of CpG sites in *RIMBP3*. Black line indicates the location of a male DMS in neutrophils (hg19 coordinates, chr22:20,459,612). (**B**) Methylation status of CpG sites in *FIGN*. Black line indicates the location of a female DMS in neutrophils (hg19 coordinates, chr2:164,590,051). E1 and E2 indicate the first and second exon of *FIGN*, respectively.

**Supplementary Figure S2.**

| **DMS type** | **CI DMS** | ***P*-value DMS** |
|---|---|---|
| CD4T_male | 63 (0) | 4,287 (274) |
| CD4T_female | 57 (0) | 3,726 (299) |
| Mono_male | 1 (0) | 2,101 (136) |
| Mono_female | 5 (0) | 1,916 (139) |
| Neu_male | 22 (1) | 2,170 (117) |
| Neu_female | 106 (5) | 3,662 (224) |
| All | 213 (6) | 15,611 (1,066) |

**Supplementary Figure S2.** Overlaps between the DMSs and methylation array platform. Numbers of DMSs identified by the CI- or *P*-value-based approach, with overlaps in parentheses based on the Infinium MethylationEPIC v1.0 B5 Manifest File (Illumina). DMS type indicates the stratification of DMSs by cell type and sex-associated methylation: CD4T_male/female, DMSs with male or female hypermethylation in CD4T; Mono_male/female, in monocytes; Neu_male/female, in neutrophils.

**Supplementary Table S1. List of DMSs identified by *P*-value-based approach**

Identified DMSs by *P*-value-based approach ($N = 17,862$) after Bonferroni correction for multiple testing based on the number of all CpGs in three cell types ($N = 69,040,758$; significance level, *P*-value $< 7.24 \times 10^{-10}$). The total number of unique DMSs across cell types was 15,611. *Chr*, chromosome; *Start* and *End*, chromosomal location of CpG dinucleotide in hg19; *DNAm_Male* or *DNAm_Female*, the mean value of DNA methylation level in males or females (%); *Z, Z*-score used for test of the difference in population proportions; *P-value*, the *P*-value from a two-sided test for *Z*; *DMS type*, a stratification of DMSs by both cell type and sex-associated methylation (CD4T_male/female, DMSs with male or female hypermethylation in CD4T; Mono_male/female, in monocytes; Neu_male/female, in neutrophils); *DMS/DMR ID*, a group name of each DMS or DMR. The value of 0 in *P*-value column indicates *P*-value $< 2.2 \times 10^{-16}$ due to the limitations of double-precision floating-point.

**Supplementary Table S2. List of DMRs identified by *P*-value-based approach**

DMRs identified by *P*-value-based approach, including two or more DMSs ($N = 1,672$). *Chr*, chromosome; *Start* and *End*, chromosomal location of each DMR in hg19; *DNAm_Male* or *DNAm_Female*, the mean value of DNA methylation level of the included DMSs in males or females (%); *# of DMSs*, the number of the included DMSs; *Length*, the length (bp) of each DMR; *DMR type*, a stratification of DMRs by both cell type and sex-associated methylation (CD4T_male/female, DMRs with male or female hypermethylation in CD4T; Mono_male/female, in monocytes; Neu_male/female, in neutrophils); *DMR ID*, a group name of each DMR.

**Supplementary Table S3. List of DMSs identified by CI-based approach**

Identified DMSs by CI-based approach ($N = 254$). *Chr*, chromosome; *Start* and *End*, chromosomal location of CpG dinucleotide in hg19; *DNAm_Male* or *DNAm_Female*, the mean value of DNA methylation level in males or females (%); *Z*, *Z*-score used for test of the difference in population proportions; *P-value*, the *P*-value from a two-sided test for *Z*; *Male_Lower*, the lower limit of the 95% confidence interval (CI) in males; *Male_Upper*, the upper limit of the 95% CI in males; *Female_Lower*, the lower limit of the 95% CI in females; *Female_Upper*, the upper limit of the 95% CI in females; *DMS type*, a stratification of DMSs by both cell type and sex-associated methylation (CD4T_male/female, DMSs with male or female hypermethylation in CD4T; Mono_male/female, in monocytes; Neu_male/female, in neutrophils); *DMS/DMR ID*, a group name of each DMS or DMR. The value of 0 in *P*-value column indicates *P*-value $< 2.2 \times 10^{-16}$ due to the limitations of double-precision floating-point.

**Supplementary Table S4. List of DMRs identified by CI-based approach**

DMRs identified by CI-based approach ($N = 22$). *Chr*, chromosome; *Start* and *End*, chromosomal location of each DMR in hg19; *DNAm_Male* or *DNAm_Female*, the mean value of DNA methylation level of the included DMSs in males or females (%); *# of DMSs*, the number of the included DMSs; *Length*, the length (bp) of each DMR; *DMR type*, a stratification of DMRs by both cell type and sex-associated methylation (CD4T_male/female, DMRs with male or female hypermethylation in CD4T; Mono_male/female, in monocytes; Neu_male/female, in neutrophils); *DMR ID*, a group name of each DMR.

**Supplementary Table S5. GO analysis of DMS-associated genes identified by *P*-value-based approach**

*DMS type*, a stratification of DMSs by both cell type and sex-associated methylation (CD4T_male/female, DMSs with male or female hypermethylation in CD4T; Mono_male/female, in monocytes; Neu_male/female, in neutrophils); *Category* shows the GO category; *Term* lists the GO term; *Genes* represents the number of DMS-associated genes in each GO term; *P-value* gives the nominal *P*-value from ToppGene; *Adjusted P* denotes the adjusted *P*-value with Bonferroni correction. MF, molecular function; CC, cellular component; BP, biological process.

**Supplementary Table S6. GO analysis with gene length correction**

*DMS type*, a stratification of DMSs by both cell type and sex-associated methylation (CD4T_male/female, DMSs with male or female hypermethylation in CD4T; Mono_male/female, in monocytes; Neu_male/female, in neutrophils); *Category* shows the GO category; *Term* lists the GO term; *P-value* gives the nominal *P*-value; *Adjusted P* denotes the adjusted *P*-value with Bonferroni correction. MF, molecular function; CC, cellular component; BP, biological process.

**Supplementary Table S7. Enrichment analysis of DMRs using GWAS SNPs**

*ID* indicates the GWAS accession number in the GWAS Catalog; *Trait*, the reported phenotype associated with the GWAS regions; *SNP*, the number of reported SNPs; *DMR type*, a stratification of DMRs by both cell type and sex-associated methylation (CD4T_male/female, DMRs with male or female hypermethylation in CD4T;

Mono_male/female, in monocytes; Neu_male/female, in neutrophils); *DMR*, the number of overlapping DMRs; *b*, the number of overlapping CpG sites in iMETHYL; *c*, the number of non-overlapping DMRs; *d*, the number of non-overlapping CpG sites in iMETHYL; *OR*, odds ratio; *P-value*, significance calculated by Fisher's exact test; *FDR* denotes the false discovery rate calculated using the Benjamini-Hochberg method (337 × 6 tests).

**Supplementary Table S8. Enrichment of the TF-binding sites in DMSs identified by CI-based approach**

Enriched TFs after Bonferroni correction for multiple testing (7,397 entries × 6 tests). *ID* indicates the experimental identifier in ChiP-Atlas; *Antigen*, the target TF in each ChIP-Seq experiment; *Cell*, the sample cell type of the ChIP-Seq experiment; *logP*, log *P*-values in Chip-Atlas; *logQ*, log *Q*-values of peak-call significance; *FE*, fold enrichment; *DMS type*, a stratification of DMSs by both cell type and sex-associated methylation (CD4T_male/female, DMSs with male or female hypermethylation in CD4T; Mono_male/female, in monocytes; Neu_male/female, in neutrophils).

**Supplementary Table S9. Results of protein-protein interaction (PPI) analysis**

*Category*, a classification of PPI term; *Term*, a name of each PPI term; *P-value*, *P*-value for enrichment; *FDR*, the false discovery rate calculated using the Benjamini-Hochberg method. The sheet name indicates a stratification of DMSs by both cell type and sex-associated methylation (CD4T_male/female, DMSs with male or female hypermethylation in CD4T; Mono_male/female, in monocytes; Neu_male/female, in neutrophils).
